# Dynamics and impacts of transposable element proliferation during the *Drosophila nasuta* species group radiation

**DOI:** 10.1101/2021.08.12.456169

**Authors:** Kevin H.-C. Wei, Dat Mai, Kamalakar Chatla, Doris Bachtrog

## Abstract

Transposable element (TE) mobilization is a constant threat to genome integrity. Eukaryotic organisms have evolved robust defensive mechanisms to suppress their activity, yet TEs can escape suppression and proliferate, creating strong selective pressure for host defense to adapt. This genomic conflict fuels a never-ending arms race that drives the rapid evolution of TEs and recurrent positive selection of genes involved in host defense; the latter has been shown to contribute to postzygotic hybrid incompatibility. However, how TE proliferation impacts genome and regulatory divergence remains poorly understood. Here, we report the highly complete and contiguous (N50=33.8Mb - 38.0Mb) genome assemblies of seven closely-related *Drosophila* species that belong to the *nasuta* species group - a poorly studied group of flies that radiated in the last 2 million years. We constructed a high quality de novo TE library and gathered germline RNA-seq data, which allowed us to comprehensively annotate and compare insertion patterns between the species, and infer the evolutionary forces controlling their spread. We find a strong negative association between TE insertion frequency and expression of genes nearby; this likely reflects survivor-bias from reduced fitness impact of TE inserting near lowly expressed, non-essential genes, with limited TE-induced epigenetic silencing. Phylogenetic analyses of insertions of 147 TE families reveal that 53% of them show recent amplification in at least one species. The most highly amplified TE is an non-autonomous DNA element DINE which has gone through multiple bouts of expansions with thousands of full length copies littered throughout each genome. Across all TEs, we find that TEs expansions are significantly associated with high expression in the expanded species consistent with suppression escape. Altogether, our results shed light on the heterogenous and context-dependent nature in which TEs affect gene regulation and the dynamics of rampant TE proliferation amidst a recently radiated species group.

## Introduction

Eukaryotic genomes are littered with transposable elements (TEs). TEs are selfish genetic elements that self-replicate via copy and paste or cut and paste mechanisms. Despite their abundance and ubiquity in genomes (Kidwell 2002), they can be highly deleterious especially when active. When they transpose, TEs can create double strand breaks and disrupt reading frames when inserted into genes (Hedges and Deininger 2007). Even when transpositionally inactive, they can induce non-allelic exchange due to sequence homology which can create devastating genome rearrangements (Athma and Peterson 1991; Kidwell and Holyoake 2001; Xiao et al. 2000; Zhang et al. 2011).

To combat their deleterious activity, eukaryotic genomes have evolved intricate defense pathways to inactivate TEs both transcriptionally and post-transcriptionally (for review see Ozata et al. 2019). Post-transcriptional silencing generally involves small RNA-targeted degradation of TE transcripts (for reviews see Czech et al. 2018; Ozata et al. 2019; Wang and Lin 2021). Transcriptional inactivation is achieved through compaction of the chromatin environment into a dense and inaccessible state, known as heterochromatin (for reviews see Richards and Elgin 2002; Elgin and Reuter 2013). This involves di- and tri-methylation to the histone H3 tail at the 9th lysine (H3K9me2/3), which in turn recruits neighboring histones to be methylated allowing heterochromatin to spread across broad domains (Nakayama et al. 2001; Lachner et al. 2001; Bannister et al. 2001; Hall et al. 2002). Interestingly, this spreading mechanism can also have the unintended effect of silencing genes nearby TE insertions (Choi and Lee 2020). Therefore, in addition to disrupting coding sequences, TE insertions can further impair gene function by disrupting gene expression (Hollister and Gaut 2009; Lee 2015; Lee and Karpen 2017).

However, even with strong repressive mechanisms, defense against TEs appears to be an uphill battle. TEs are among the most rapidly changing components of eukaryotic genomes. TE content can differ drastically even between closely related species and has been shown to be a key contributor to genome size disparities. In *Drosophila*, the P-element, a DNA transposon originating from *D. willistoni*, invaded both *D. melanogaster* (Anxolabéhère et al. 1988; Daniels et al. 1990) and subsequently *D. simulans* (Kofler et al. 2015). Both of these cross-species invasions occurred rapidly within the last century and resulted in world-wide sweeps of the P-element in wild populations. Mobilization events are accompanied by reduction in host fertility and viability (Kidwell et al. 1977; Kidwell and Novy 1979; Schaefer et al. 1979), which in turn creates strong selective pressure for the host to evolve an updated repressive mechanism(Simkin et al. 2013; Kelleher and Barbash 2013). Such dynamics create an evolutionary arms-race between host suppression mechanisms and TE suppression escape, and is thought to underlie the recurrent adaptive evolution of many proteins involved in the TE silencing pathways (Parhad and Theurkauf 2019; Luo et al. 2020). Rapid evolution of TEs and the repressive pathways have even been implicated in establishing postzygotic reproductive isolation between closely related *Drosophila* species (Kliman et al. 2000; Garrigan et al. 2012; Brand et al. 2013).

Beyond their deleterious potential, TEs can also be sources of novelty in the genome (Kidwell and Lisch 1997). TEs, or parts of their sequences, have been co-opted for gene regulatory functions such as promoters and enhancers (Jacques et al. 2013; Merenciano et al. 2016; Sundaram and Wysocka 2020). Their recurrent transpositions across nascent sex chromosomes also mediated the evolution of dosage compensation chromosome-wide (Ellison and Bachtrog 2013; Zhou et al. 2013). Insertions of TEs to the proximity of genes have also been shown to create functional chimeric retrogenes (Buzdin 2004; Xing et al. 2006). In mammals, KRAB-zinc finger transcription factors have repeatedly co-opted the transposase protein encoded by DNA transposons, allowing for the diversification of their binding targets (Cosby et al. 2021). Lastly, in flies, domesticated retrotransposons insert at chromosome ends for telomere extension thus alleviating the need for telomerase to solve the end-replication problem (Traverse and Pardue 1988; Biessmann et al. 1990; Levis 1994). Therefore, TEs do not just force the host defense to adapt in order to suppress their activity, but they can also be beneficial drivers of genome evolution (Kidwell and Lisch 1997; Casacuberta and González 2013).

While TEs can have multi-faceted influences on the genome and its evolution, the dynamics of TE amplification and suppression escape remain poorly understood, especially outside of select model species. This is in part due to the inherent challenge associated with studying highly repetitive sequences, an issue that became particularly problematic during the boom of short-read sequencing technologies in the last two decades. Most TE-derived short reads (typically less than 150bps) cannot be uniquely assigned to a region of the genome, which causes errors in mapping and breakages in genome assemblies (Bourque et al. 2018; O’Neill et al. 2020). Numerous approaches have been devised that take advantage of different features of short-read sequencing platforms (e.g. paired sequencing) to call insertions (Linheiro and Bergman 2012; Cridland et al. 2013; Rahman et al. 2015; McGurk and Barbash 2018; Wei et al. 2020), but such methods are nevertheless limited by short read lengths, often producing inconsistent results (Vendrell-Mir et al. 2019). With the advent of long-read (5kb+) sequencing technologies from Oxford Nanopore and PacBio, many of these issues can finally be circumvented (Hotaling et al. 2021). The use of such technologies have already led to drastic improvements of genome assemblies across highly repetitive genomes in, for example, flies (Mahajan et al. 2018; Bracewell et al. 2019; Chakraborty et al. 2021), mosquitoes (Matthews et al. 2018), mammals (Bickhart et al. 2017), and humans (Nurk et al. 2021)..

Highly contiguous genomes with well-represented repeat content permit comprehensive analyses of TE insertions across the genome. Multiple such high quality genomes further enable analyses of the dynamics of TE proliferation through a comparative and phylogenomics framework. Therefore, to illuminate how TEs proliferate and potentially drive genome evolution and speciation, we used long-read technologies to generate high quality genome assemblies of seven closely related *Drosophila* species (**Figure 1A,B**) that belong to the *nasuta* group. These species group radiated in the last two million years (Kitagawa et al. 1982; Bachtrog 2006; Ranjini and Ramachandra 2013; Mai et al. 2020) and is widely distributed across Asia, with some populations found in eastern Africa, Oceania, and Hawaii (Wilson et al. 1969; Mai et al. 2020), and *D. nasuta* has recently been identified as an invasive species in Brazil that is spreading quickly in South America (Vilela and Goñib 2015; Silva et al. 2020). While most of the species are geographically isolated, they have varying levels of reproductive isolation (Kitagawa et al. 1982); over half of interspecific crosses produce viable offspring. With these high quality genomes, we sought to systematically understand how TE insertions around genes affect gene expression, and how frequently TEs escape repression and expand. To answer these questions, we generated a library of a common set of high quality TE consensus sequences from de novo TE calls across the genome assemblies. With this library, we identified species-specific TE insertions and found that TEs frequently expand, likely due to suppression escape, with >50% of TEs showing evidence of lineage-specific expansion in at least one species. Species-specific TEs are disproportionately found near lowly expressing genes and rarely have impact on gene expression. Lastly, we show that silencing of expanding TEs can lead to silencing of neighboring genes.

**Figure 1.**
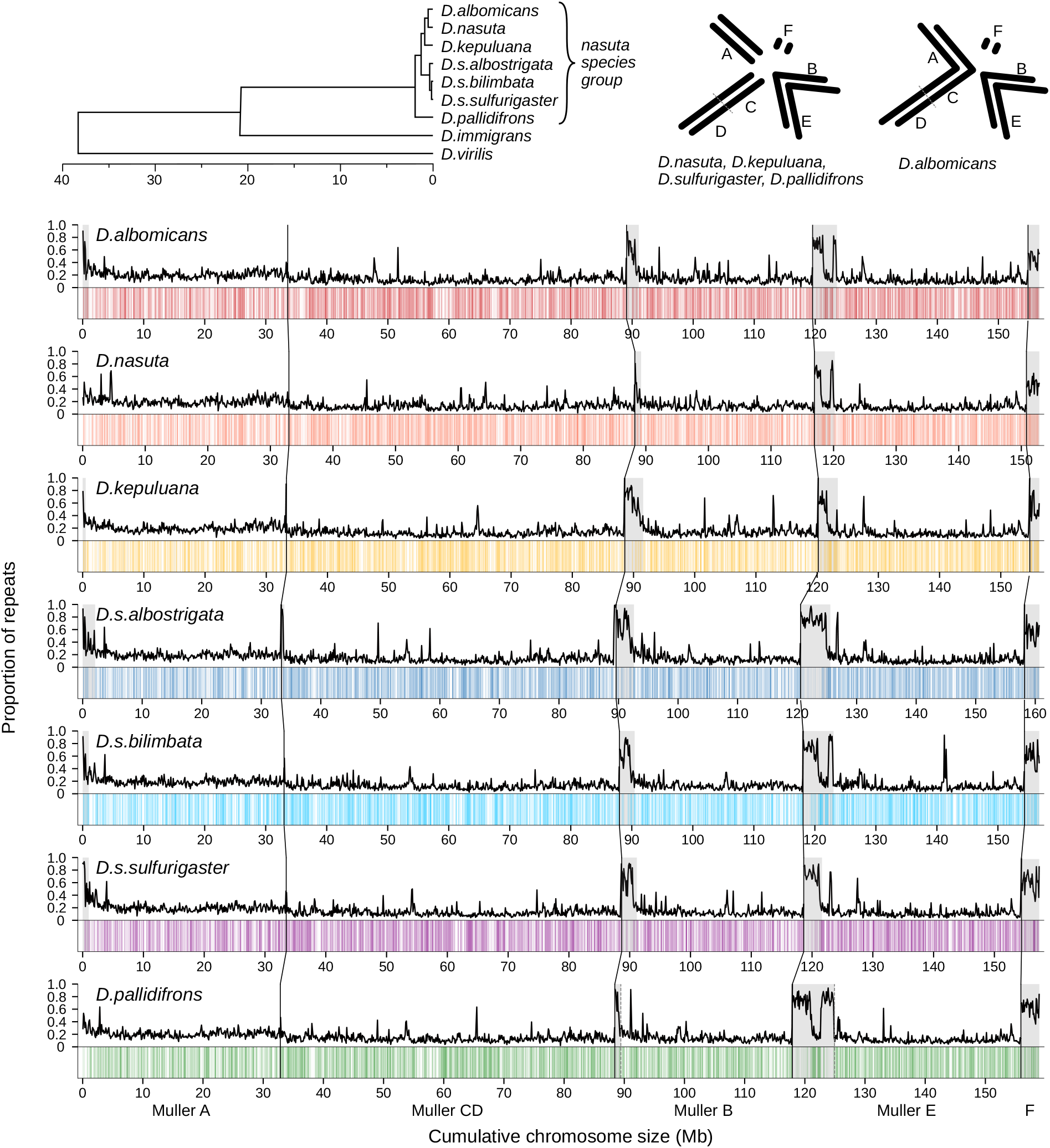
Genomes of the Drosophila nasuta species group. A. Phylogeny of the *nasuta* species radiation within the *Drosophila* subgenus. Tree adapted from (Mai et al. 2020) and (Izumitani et al. 2016) B. Karyotypes of the species group; chromosomes are oriented such that centromeres are pointed towards the center of circle. C. Long read-based genome assemblies of seven species. For each species, the top track depicts the repeat content estimated for 100kb windows. Positions of annotate genes are represented on the bottom track as vertical lines. The centromeric end are on the left side of each chromosome. Regions deemed as pericentromeric are highlighted in gray. Chromosomes are demarcated by black vertical lines. Unless otherwise stated, species are represented by colors used here: red (*D. albomicans*), orange (*D. nasuta*), yellow (*D. kepuluana*), navy (*D. s. albostrigata*), purple (*D. s. bilimbata*), purple (*D. s. sulfurigaster*), and green (*D. pallidifrons*).

## Results

### High quality genome assemblies across seven species

Genome assemblies for females of seven species in the *nasuta* clade—*D. albomicans*, *D. nasuta*, *D. kepulauana*, *D. sulfurigaster albostrigata*, *D. sulfurigaster bilimbata*, *D. sulfurigaster sulfurigaster*, and *D. pallidifrons*—were generated using Nanopore and Hi-C reads (**Table 1**; **Figs. S1-S7**). The methodology for preparing reads was adopted from Bracewell et al. and applied across all species: error-correct Nanopore reads with canu, generate contig assembly with wtdbg2 and flye, polish assembly with racon and pilon, remove contigs that belong to other organisms with BLAST, and stitch contig assemblies using Hi-C reads as input for Juicer and 3d-dna (Bracewell et al. 2019; Koren et al. 2017; Ruan and Li 2020; Kolmogorov et al. 2019; Vaser et al. 2017; Walker et al. 2014; Durand et al. 2016; Dudchenko et al. 2017; Altschul et al. 1990). However, there is no universal pipeline to generate ideal assemblies; the assembly pipeline for different flies underwent various adjustments for optimal results (see Methods). Overall, we generated consistent assemblies for each species using an average of 30.3x long read coverage (std dev = 10; **Table S1**), resulting in a mean N50 of 35.9 Mb (std dev = 1.4 Mb; **Table 1**), assembly size of 166.6 Mb (std dev = 2.59 Mb; **Table 1**), and BUSCO score of 99.3% (std dev = 0.4%; **Table S2**).

**Table 1.**
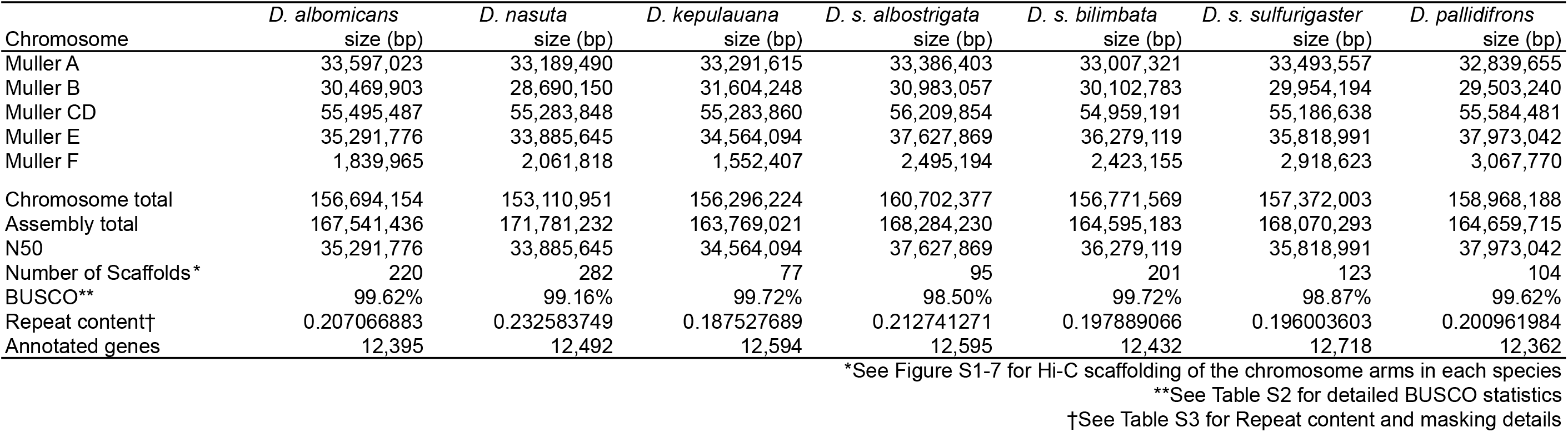
Size of genome assemblies (and their chromosomes) for each species and their associated summary statistics.

We leveraged the chromosome level genome assemblies alongside RNA-seq data from *D. albomicans* and *D. nasuta* (Zhou and Bachtrog 2012) to annotate genes across all species (**Table 1**). An average of 12,513 genes were annotated per species (std dev = 128.48), which is lower than the number of genes annotated in other *Drosophila* species (Consortium and Drosophila 12 Genomes Consortium 2007). In order to analyze homologous genes, we clustered genes between species with OrthoDB and found 9,413 genes shared across all species (Kriventseva et al. 2019).

### Generating a curated de novo TE library

For each genome, we used RepeatModeler2 to generate a de novo TE library, which we then used to annotate the genome (Flynn et al. 2020). This resulted in between 18.8%-23.3% of the genomes being masked (**Table S3**). Further, high repeat content near chromosome ends show that these near-chromosome length scaffolds include some heterochromatin and pericentromeric regions. Expectedly, gene density and repeat density are negatively correlated (**Figure 1C**).

One major challenge with de novo TE identification using standard computational methods is that the resulting TE libraries are littered with redundant and fragmented entries. Furthermore, we find that secondary structures such as nested insertions or fragment duplications (**Figure S8**) are frequently identified as unique TE entries in the libraries. To improve the de novo TE library and to generate a common set of TE consensus sequences across all the *nasuta* subgroup, we devised a pipeline that utilizes multiple steps and metrics (**Figure 2A**). After an initial de novo TE library call with RepeatModeler2 for each of the genomes, we demarcated the euchromatin/pericentromere boundaries (**Figure 1C**). Reasoning that recently active TEs are more likely to be intact and surrounded by unique sequences in the euchromatin, we then ran RepeatModeler2 for a second time on only the euchromatic portions of the genome assemblies. The resulting TE libraries were then merged across all the species generating a library of 1818 entries.

**Figure 2.**
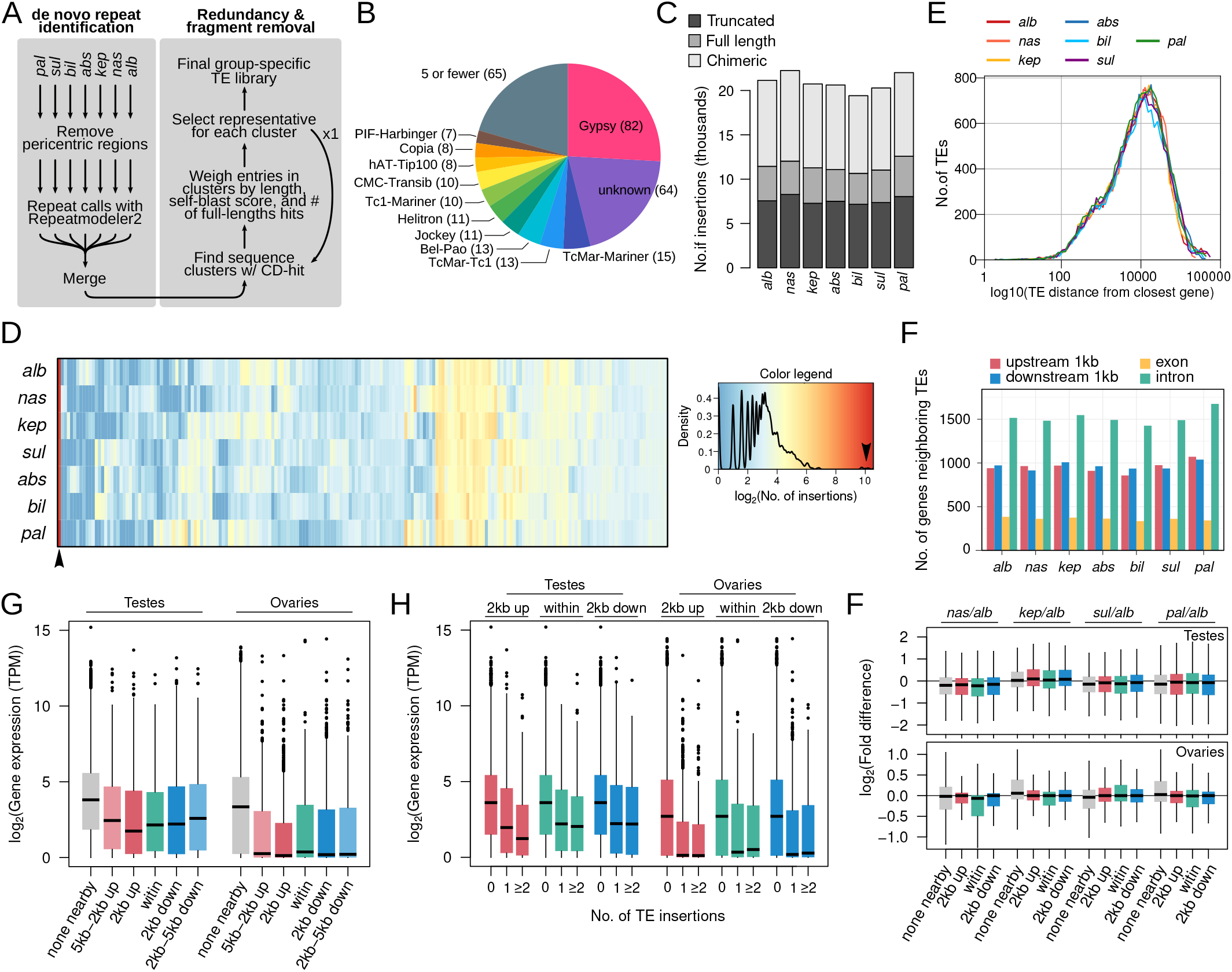
De novo identification and distribution of TE insertions across the genomes. A. Pipeline to construct and refine de novo TE reference from genome assemblies. We used RepeatModeler2 to first identify repeats from the euchromatic regions of each species. The resulting repeat libraries are merged followed by sequence clustering with CD-HIT. Multiple indexes were used to select the full length representative TEs. B. Breakdown of TE classes identified; for breakdown of the grey section see supplementary figure S9. C. Number of full length and truncated insertions found in each genome. The chimeric class represents the merger of annotations that overlap or are contiguous. D. Copy number of full length insertions of 318 TE families across the seven genomes. E. Distribution of the distance between TEs and genes across the species. Intergenic insertions are not counted. See supplementary figure S9 for distribution of intergenic insertions from exons . F. Number of genes with TEs inserted in different regions of genes with and without insertions. G. Transcript abundance of annotated genes in Transcripts per million (TPM), subsetted into different classes depending on where TE insertions are found. H. Transcript abundance of genes with different numbers of TE insertions. I. Fold-difference in transcript abundance of orthologous genes depending on different numbers of insertions in *D. albomicans*. See supplementary figure S10, for comparisons using insertions in other species.

We then used CD-HIT2 to group the entries into clusters based on sequence similarity (Fu et al. 2012). By default, CD-HIT2 outputs the longest sequence in each cluster. While this means full length entries will be favored over fragmented entries (when both exist in the library), entries with nested structures or chimeric TEs will be selected in favor of full-length but shorter elements. Therefore, in addition to sequence length, we evaluate each TE in each cluster based on two additional metrics to preferentially select representative and full length TE consensus sequences. While increasing entry lengths, chimeric TEs are unlikely to be frequently found in the genome; we therefore blasted the TE entries to the genome and tallied the number of times hits cover 80% of the length of the entries. In addition, we blasted the TEs to themselves to determine internal redundancy; entries with internal duplications or nested insertions will have a high self-blast score. We then selected the representative sequence as the longest sequence with high number of near full length blast hits and low self-blast score. We then repeated this step one more time to further remove redundancies in the library. After these two rounds of clustering with CD-HIT2, the TE library size was reduced to 351 consensus sequences. Afterwards, we merged TE sequences that make up a larger, full length element through patterns of co-occurrences in the genome. This resulted in a substantially reduced library with 318 entries.

The TEs generated from RepeatModeler2 are, by default, assigned to a TE category. To validate these assignments, we used ClassifyTE to reannotate the TE library (Panta et al. 2021). There is a 50% concordance between the annotations from RepeatModeler2 and ClassifyTE. Entries that were different between the two annotations were assigned the default category from RepeatModeler2. Gypsy elements make up the majority of the TE library, consisting of 82 entries (25.8%) followed by unknown families (57 entries, 17.9%; **Figure 2B**). All other TE families make up less than 5% of the TE library. The pattern of high number of Gypsy families is similar to those in other *Drosophila* species (Mérel et al. 2020).

### TE Insertion patterns across the genome

Using the refined *nasuta* group-specific TE library, we annotated TE insertions in each genome assembly using RepeatMasker (see Materials and methods). We classified full length insertions as annotations that cover at least 80% of the entry in the library; insertions covering less than 80% and are over 200bp are classified as truncated insertions. In addition, we merged annotations that are contiguous or overlapping, which can be due to nested insertions or remaining redundancies in the repeat library. The number of full length TEs range from 3489 to 4544 (**Figure 2C**) and the majority (73.6% on average) fall within euchromatic regions of the genomes, (**Figure S9C**), similar to previous reports (Biémont and Vieira 2005; Consortium and Drosophila 12 Genomes Consortium 2007). Truncated insertions are nearly 2x as numerous (ranging between 7164-8273; **Figure 2C**). As expected given their mosaic nature, the merged annotations have the largest fractions fall within the heterochromatic regions (**Figure S9**). With the exception of four TE families, all are found in low to intermediate copy numbers with fewer than 100 copies in any given genome, consistent with previous findings (**Figure 2D**). Interestingly, one TE stands out as having thousands of copies across all the genomes (**Figure 2D**, arrowhead, see section below).

To evaluate if and how TEs impact gene function, we looked at TE insertion patterns with respect to neighboring genes (**Figure 2E**, **Figure S9A**). On average, TE insertions are 18.5 kb away from the nearest genes; 41.5-46.9% of insertions are within 5kb of genes (**Figure 2E**). Of the 12,362-12,718 genes annotated, 2,887 to 3,343 have insertions within or nearby (<1kb 5’ or 3’). Of those, 48.9% to 50.1% of genes have insertions within introns, which would not affect the reading frame (**Figure 2G**).

### TE insertions are associated with low expression of nearby genes

To systematically examine the impact, if any, of TE insertions on gene expression, we generated ovarian and testes mRNA-seq for five of the seven species investigated (excluding *D. s. bilimbata* and *D. s. albostrigata)*. Genes with TE insertions nearby or within are over-represented for lowly expressed genes, in both testis and ovaries (p < 2.2e-16 Wilcoxon rank sum test; **Figure 2G**; **Figure S10**). Genes with insertions less than 2kb upstream have the lowest expression in both the testes and ovaries (**Figure 2G**). Genes with TEs inserted further away (2-5kb) also have significantly lower expression, though to a lesser extent (**Figure 2G**, **Figure S10**). Moreover, we find that gene expression is inversely correlated with the number of TE insertions (**Figure 2H**). This negative relationship holds for insertions found within, upstream, and downstream of genes. Interestingly, ovarian expression appears to be more negatively associated with TE insertions, with no expression in ovaries of nearly half of the genes with TEs inserted nearby (**Figure 2G, H**).

Due to the spreading of heterochromatin, TE insertions can induce epigenetic silencing at neighboring genes (Choi and Lee 2020). Therefore, prima facie, these results are consistent with the epigenetic silencing of genes due to neighboring TE insertion. To further test this, we reasoned that if TE insertions are inducing downregulation of surrounding genes, orthologous genes without insertions should be more highly expressed. To test this possibility, we compared the expression of orthologs when insertions are found in one species but not the other. Curiously, we do not find that expression between orthologs changes significantly depending on the presence of insertions nearby or within (**Figure 2F**, **Figure S11**). This suggests that insertions within/nearby genes are not systematically downregulating expression. Instead, TEs appear to preferentially insert and/or accumulate around lowly expressed genes.

### Survivor bias likely drives anti-correlation between TE insertion and gene expression

To elucidate the source of the negative association between gene expression and TE insertions nearby, we looked at all TE insertions found around/within the 9413 genes with orthologs across all species. To ensure that only unique insertions are counted and ancestral insertions are counted only once, we removed insertions belonging to the same TE family that are within 100bp relative to the neighboring genes. Further, we removed pericentric genes from these analyses to avoid their high local TE counts driving correlations. For these gene orthologs, we indeed find a significant negative correlation between TE insertion counts and averaged gene expression in both testes and ovaries (**Figure 3A, B**; p < 2.2e-16). Similar correlations are also found when looking at the proportion of bases covered by TEs around and within genes (**Figure S12**). Curiously, the negative correlation of ovarian expression is significantly stronger than that of testes expression (**Figure 3A, B**; p < 1e-8, Pearson and Filon’s z).

**Figure 3.**
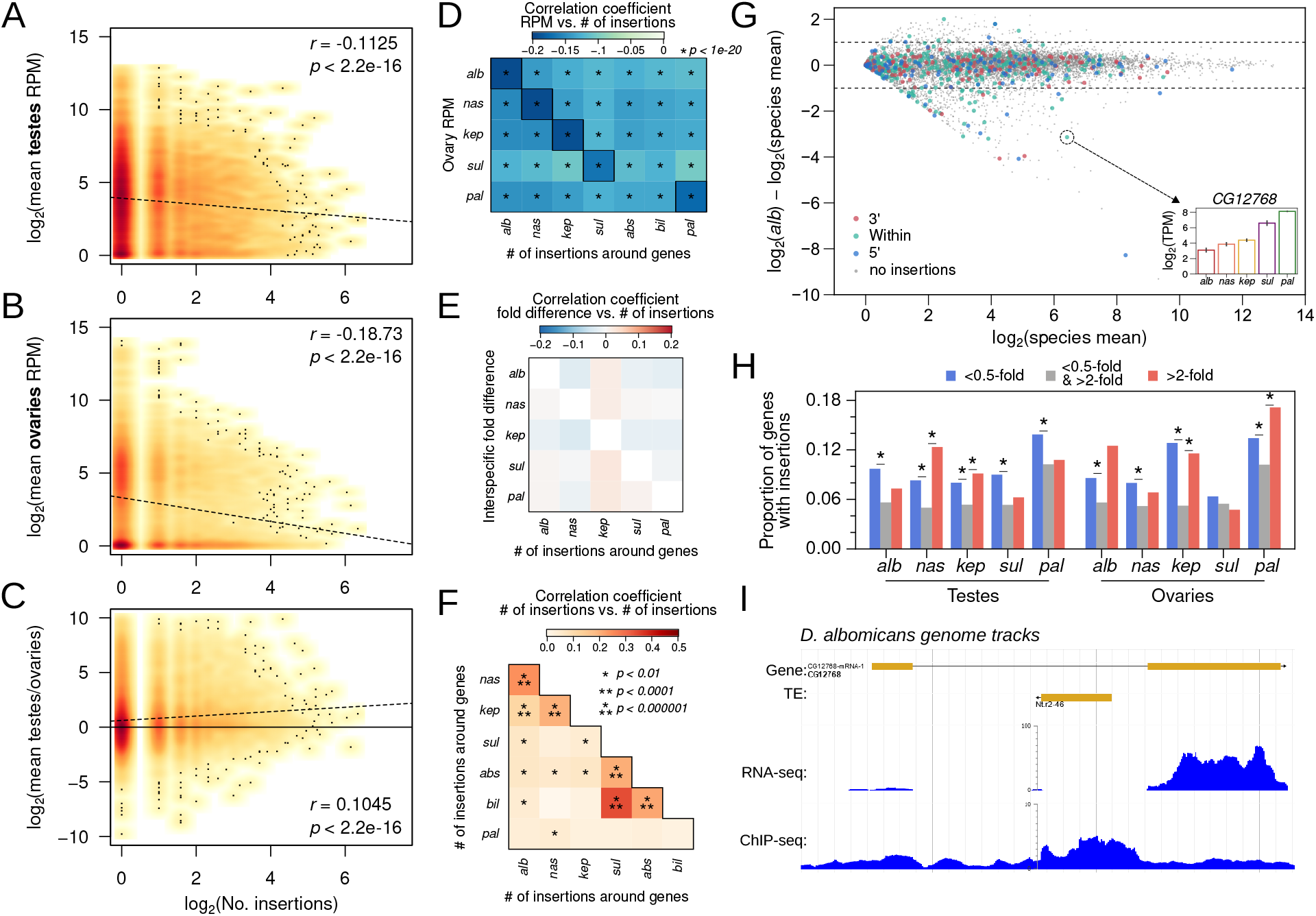
Negative association between TE insertions and genic expression. A-B. Density scatterplots of number of unique (both full length and truncated) TE insertions around genes (±2kb) across all the *nasuta* species genomes plotted against genic transcript abundances (averaged across the species) in the ovaries (A) and testes (B). Increased intensity of warm colors indicate higher density of points. Scattered black dots indicate positions of single points. Regression lines are depicted by dotted lines; the Pearson’s correlation coefficients and corresponding p-values are labeled in the top right. C. Same as A and B, but with the fold difference of genic expression between testes and ovaries. D. Pairwise correlation of TE insertion counts around genes in a particular species to the ovarian transcript abundance of the gene orthologs in another species. E. Pairwise correlation of TE insertion counts around orthologous genes across species; genes with no insertions in either species are not used. G. MA-plot of average gene expression (TPM) across species in the testes (x-axis) plotted against fold difference between the D. albomicans expression and the average across species (y-axis). Colored points represent genes with TE insertions in different parts of the gene. Horizontal dotted line demarcates 0.5- and 2-fold differences. Inset shows the testes expression of the CG12768 across all five species. For MA-plots in ovaries and other species, see supplementary figure 13. H. Proportions of genes with TE insertions with low and high gene expression relative the the species average (i.e genes below or above the dotted lines in A), for each species and in ovaries and testes. I. Genome browser shot of CG12768 showing tracks for gene structure, TE insertions, transcript abundance, and H3K9me3 enrichment. For genome browser shot of this gene in other species see supplementary figure.

We then looked at the extent of correlation between species-specific TE insertion counts to gene expression across species. If TEs insert independently at different genes and are down-regulating nearby genes in one species, we expect no cross-species correlations. Instead, significantly negative correlations are observed between all pairwise comparisons (**Figure 3D**), although the within-species correlations are significantly more negative than between-species correlations (**Figure 3D**, outlined boxes). Further, insertion-induced epigenetic down-regulation to neighboring genes is expected to increase expression divergence between species, since genes with insertions are expected to be more lowly expressed than their orthologs without insertions. We do not find any significant correlation between insertion counts around genes in one species and their expression fold-differences when compared to orthologs without insertions (**Figure 3E**). However, when comparing the distribution of TEs between species, we find that the number of TE insertions at/near genes are correlated between many of the species (**Figure 3F**). Especially between more closely related species pairs, the correlation of insertions are highly significant, suggesting that TEs have a tendency to independently insert and/or accumulate near the same genes in different genomes. Thus, between-species correlations in TE counts vs. gene expression (**Figure 3D**), and low interspecific expression divergence (**Figure 3E**) may in part be explained by the same genes being targeted by TEs in different species. Biased insertion counts near lowly expressed genes could be due to insertion bias or survival bias. The former can result from TEs preferentially targeting specific genomic features to insert such as promoters and accessible chromatin; the latter is likely the result of low fitness consequences due to insertions near lowly expressed genes.

### TE insertions associated with extreme expression changes in a small number of genes

TE insertions do not appear to have pervasive silencing effects on neighboring genes (**Figures 2G**, **3B-C**, **4D**). However, there are known cases where individual TE insertions modulate gene regulation of nearby genes. To identify such cases, we compared the expression of each gene in each species to the average expression across all species (**Figure 3G**, **Figure S13**). For the vast majority of genes with/nearby insertions, their expression does not deviate from the cross-species average. However, interestingly, we notice multiple cases where insertions are associated with substantially lowered gene expression. Examining the small fraction of genes with expression less than half of the cross-species average, we find that there are between 55-167 genes in each species showing low expression and nearby/intronic insertions (**Table S4**). Consistent with TE-induced epigenetic silencing, these genes with reduced expression are significantly overrepresented for genes with TE insertions in almost every species, and in both ovaries and testes (**Figure 3H**).

To determine whether TE insertions are inducing epigenetic silencing of nearby genes in some of these cases, we selected on one of the more significantly downregulated genes, *CG12768*, which has an insertion in the first intron (**Figure 3I**) and shows the lowest expression in *D. albomican* testes (**Figure 3G**, inset). Accompanying its low expression in *D. albomicans*, we find elevated enrichment of H3K9me3 at the intronic insertion as well as across the gene body, exons and 5’ region (see below for ChIP analysis). Notably, this insertion did not appear to completely silence the gene, as abundant RNA-seq reads still map to the second exon, albeit substantially lower than other species (**Figure S14**).

Interestingly, TE insertions are not just associated with highly downregulated genes: we find that highly upregulated genes in a species (>2-fold higher than species mean) can also be significantly over-represented by genes with insertions. While not significant in all species, up-regulated genes have proportionally more TE insertions in all comparisons (**Figure 3G**). For example, the gene *Gyc88E* in *D. albomicans* has an intronic insertion in the first exon and is the highest expressed orthologs in the testes (2.16-fold higher than the next highest; **Figure S15**). Therefore, TE insertions appear to be associated with increased expression divergence through both down- and up-regulation of nearby genes.

### H3K9me3 spreading around TE insertions near genes

To evaluate the extent to which epigenetic silencing of TEs can lead to reduction in expression of neighboring genes, we analyzed available ChIP-seq data for the repressive heterochromatic histone modification H3K9me3 in *D. albomicans* male 3rd instar larvae (Wei and Bachtrog 2019). We examined the extent of H3K9me3 spreading from TE insertions with different distances to the closest gene; to avoid TEs inside the pericentromeric or telomeric heterochromatin, we analyzed only those >5Mb from the chromosome ends. Insertions over 5kb from genes show the highest H3K9me3 enrichment in neighboring regions (**Figure 3A**, top). TEs that are closer to genes (within 5kb of genes), on the other hand, show lower levels of heterochromatin spreading. Less heterochromatin spreading from TE insertions nearby genes is consistent with opposing effects of heterochromatin formation and gene expression; transcriptionally active chromatin near genes may impede the spreading of silencing heterochromatin. Looking more closely, we find that high H3K9me3 enrichment is observed in the immediate vicinity up and downstream of the insertions and quickly drops off within 100bp (**Figure 4A**, bottom). Interestingly, this rapid decline from highly elevated H3K9me3 enrichment is observed regardless of insertion distance. Therefore, despite a narrower spreading range of TEs close to genes, the silencing effect in the immediate vicinity is similar to those far from genes, and may explain the paucity of insertions within 100bp of genes (**Figure 2E**) and exons (**Figure S9A**).

**Figure 4.**
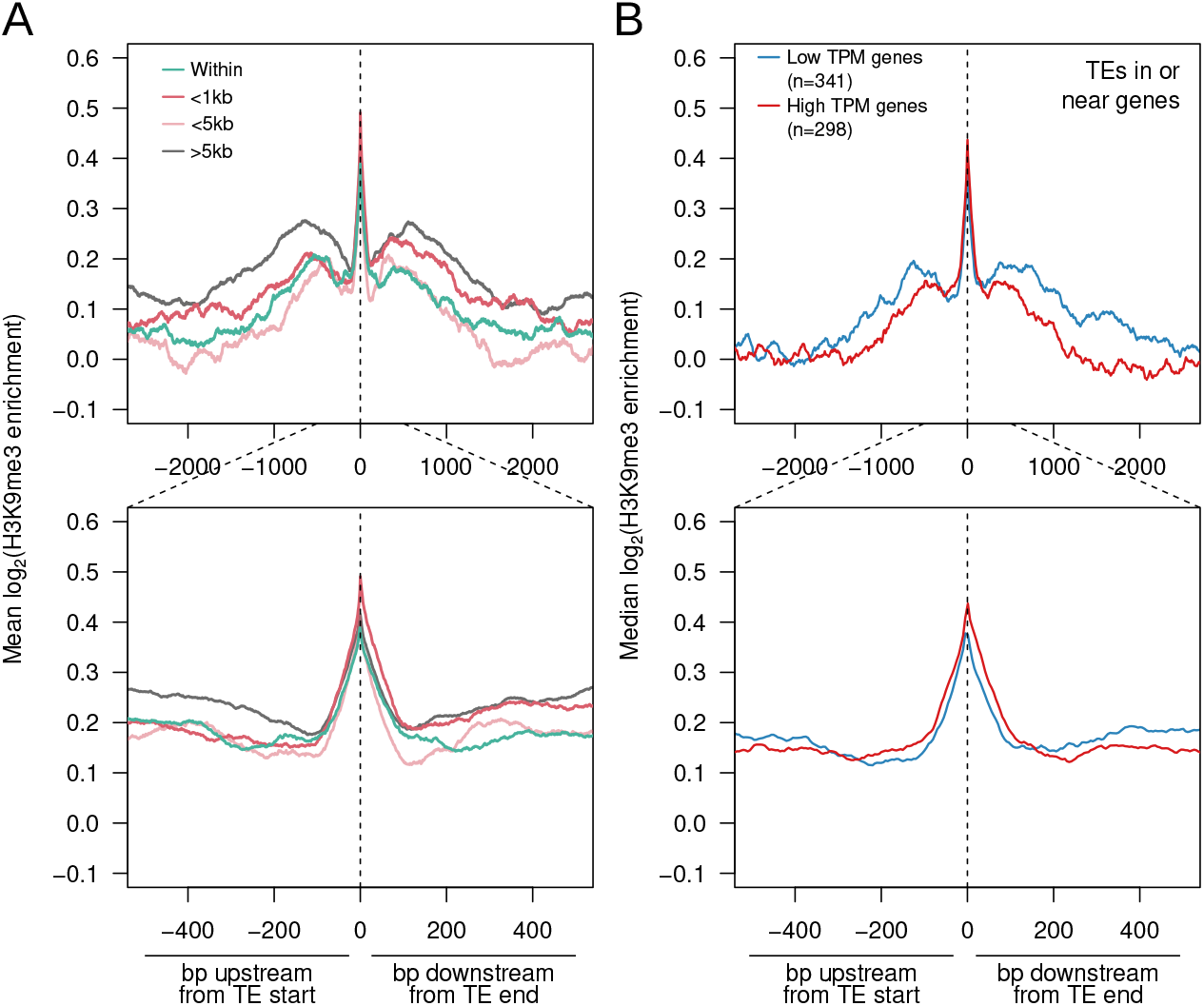
Epigenetic silencing through H3K9me3 spreading around TE insertions. A. Median H3K9me3 enrichment ± 5kb upstream and downstream of TEs inserted at different distances to genes (enrichment across TE insertions not plotted). TE insertions within pericentric regions are removed from analyses. Zoomed in plot (±500 bp) is shown below. B As with A but with TEs inserted within genes or <2kb around (C) genes of different expression levels.

To address whether heterochromatin spreading from TEs reduces expression of nearby genes, we evaluated the extent of H3K9me3 enrichment surrounding TE insertions that are nearby genes with different expression levels in testes. Insertions were partitioned by their proximity to genes with low (< 8TPM) and high expression (>8 TPM). Insertions around low TPM genes show a higher H3K9me3 enrichment and spreading than those around high TPM genes (**Figure 4B**, top). While these differences are consistent with epigenetic silencing of genes induced by neighboring TEs, they could also reflect high transcriptional activity opposing heterochromatin spreading from nearby TEs. Given the lack of systematic downregulation between genes with insertions and their orthologs (**Figure 2F**), yet overrepresentation of TE insertions in genes that are downregulated (**Figure 3H**), our data suggest that both forces are at play.

### Recurrent and rampant amplifications of DINEs

The most abundant TE, accounting for 2.1-3.8 Mb across all the species, is a 770 bp repeat which shows homology to the Drosophila INterspersed Element (DINE) - a non-autonomous DNA transposon that is highly species-specific (Locke et al. 1999). DINE’s are widespread in the *Drosophila* genus, with hundreds to thousands of copies identified across a wide range of *Drosophila* species (Yang and Barbash 2008). They appear particularly abundant in the *nasuta* species complex, with 1501-3202 full length and 4863-6793 truncated DINE insertions identified across species.

Phylogenetic analysis of individual TE insertions can reveal about their evolutionary history, including the timing of when a particular TE likely was transcriptionally active. To study the explosion of DINE elements in the *nasuta* species group, we determined their phylogenetic relationship, using near-full length copies with the addition of insertions found in the *D. immigrans* genome as the outgroup (**Figure 5A**). We find a complex phylogenetic tree where the majority of DINEs do not show species-specific clustering. Instead, insertions from different species in the *nasuta* subgroup are highly intermingled, indicating that the bulk of DINE amplification predated the radiation of this species complex (**Figure 5A**). Most of the elements are likely currently inactive given the lack of species-specific clusters and long terminal branches (**Figure S16**).

**Figure 5.**
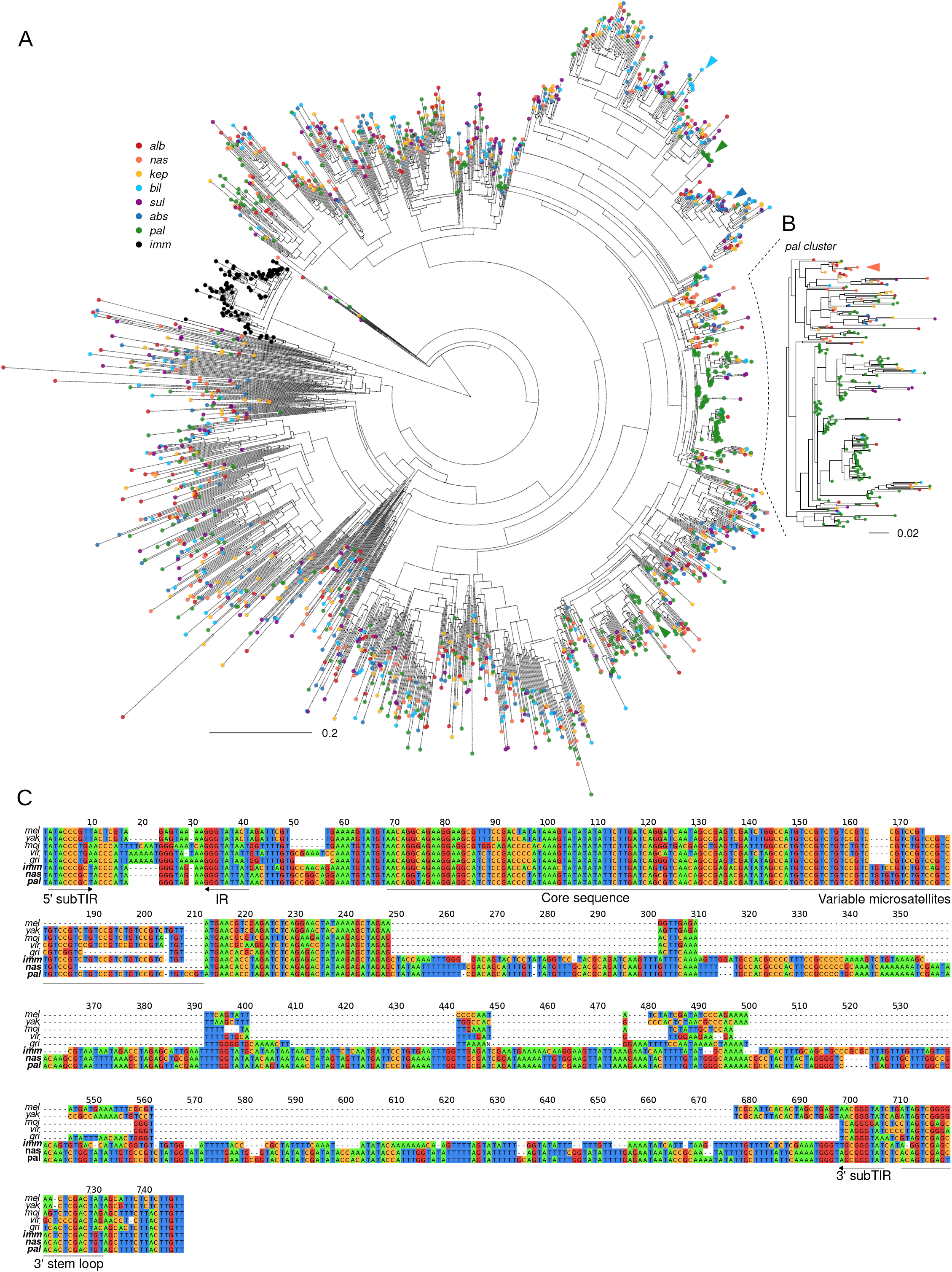
Recurrent DINE expansions. A. Radial tree of subsampled DINE insertions with the addition of *D. immigrans* DINE elements as outgroup. Insertions from the same species have the same colored tips. Colored arrowheads point to small scale species-specific expansions on the tree. B. Large cluster of *D. pallidifrons* DINE insertions indicate recent burst of species-specific activity. C. Multiple sequence alignments of consensus DINE sequences of representative species. DINE-specific sequence features are annotated beneath the tracks.

While most DINEs in the *nasuta* subgroup likely originated from old expansion events, we nevertheless identified multiple instances of species-specific clustering. First, we find that the *D. immigrans* DINEs form a monophyletic clade with short branch lengths, suggesting a relatively recent, *immigrans*-specific expansion of this element. Second, we identified multiple clusters of *D. pallidifrons* insertions throughout the tree, including one large branch containing 142 out of 400 (subsampled) DINE insertions. *D. pallidifrons* DINEs within this branch contain several distinct clusters with short branch lengths, suggesting that multiple copies of DINE are currently (or have been recently) amplifying in the genome (**Figure 5B**). Expansions of DINE in *D. pallidifrons* and *D. immigrans* are consistent with a small number of elements (if not a single copy) escaping silencing, which subsequently generated a large number of insertions. Interestingly, multiple smaller clusters of *D. pallidifrons* DINE expansion (**Figure 5**, green arrows) are also found in distant branches across the phylogeny, suggesting that other DINE lineages may have reactivated (see discussion). Lastly, though less obvious, smaller scale copy number increases of DINE can also be observed in other species, such as the large numbers of *D. albomicans*, *D. nasuta* and *D. kepulauana* DINEs within the *D. pallidifrons* cluster that suggest both species-specific insertion events as well as older insertions events in their common ancestor. Similarly, small scale expansion events are also observed for the *sulfurigaster* species complex.

To better understand the sequence changes that may have precipitated the expansions, we first generated consensus sequences for DINEs in *D. immigrans*, across the *nasuta* subgroup, and in specifically the *D. pallidifrons* cluster (**Figure 5B**) from the *D. pallidifrons* genome. We then compared them to the previously reported consensus sequences from other *Drosophila* species (**Figure 5C**). While DINEs are between 300-400bps in the other species, they double to 695 and 726 bp in *D. immigrans* and the *nasuta* group, respectively. However, they still contain many of the main features such as the presence of sub-terminal inverted repeats, microsatellite regions consisting of variable lengths of simple repeats and 3’ stem loop. Conservation can be found across the core sequence near the 5’. Nearly all the sequence length increase can be found in the middle disordered region where alignment is poor even between *D. virilis* and *D. melanogaster*. We note that there are several indels and SNPs that differentiate between consensus from the *nasuta* group consensus and the *pallidifrons* cluster. However, many of these mutations are found in DINEs that are outside of the expanded clusters.

### Frequent expansion likely due to suppression escape

Given the pattern of proliferation of the DINEs, we were curious as to the frequency in which TEs can escape suppression and expand. We therefore generate phylogenetic trees of 147 TEs where we can find more than 20 copies across all seven species; expansions were identified as branches showing significant lineage and/or species-specific clustering (**Figure 6A-D**). We find that 78 TEs show significant species-specific clustering in at least one species, suggesting TE proliferation occurs frequently in different species (**Figure 6A**). In most cases, individual TE expansions do not reach beyond 50 copies. Expansion occurs across all types of elements although in different ways (**Figure 6A**). For example, for a variant of the Gypsy LTR retrotransposon, expansions are observed in four species as well as prior to the *sulfurigaster* semi-species split (**Figure 6B**). In contrast, for Merlin, a DNA transposon, expansions are observed in *D. pallidifrons* and *D. nasuta* and prior to the *D. albomicans/D. nasuta /D. kepulauana* species split (**Figure 6C**). Lastly, a rolling circle element expanded in *D. pallidifrons* and two of the *sulfurigaster* species (**Figure 6D**). Strikingly, there are 47 expanded TE families in *D. pallidifrons* which accounts for its higher repeat content compared to the other species (**Figure 6C-E**) and may suggest increased tolerance to TE load and/or reduced genomic defense.

**Figure 6.**
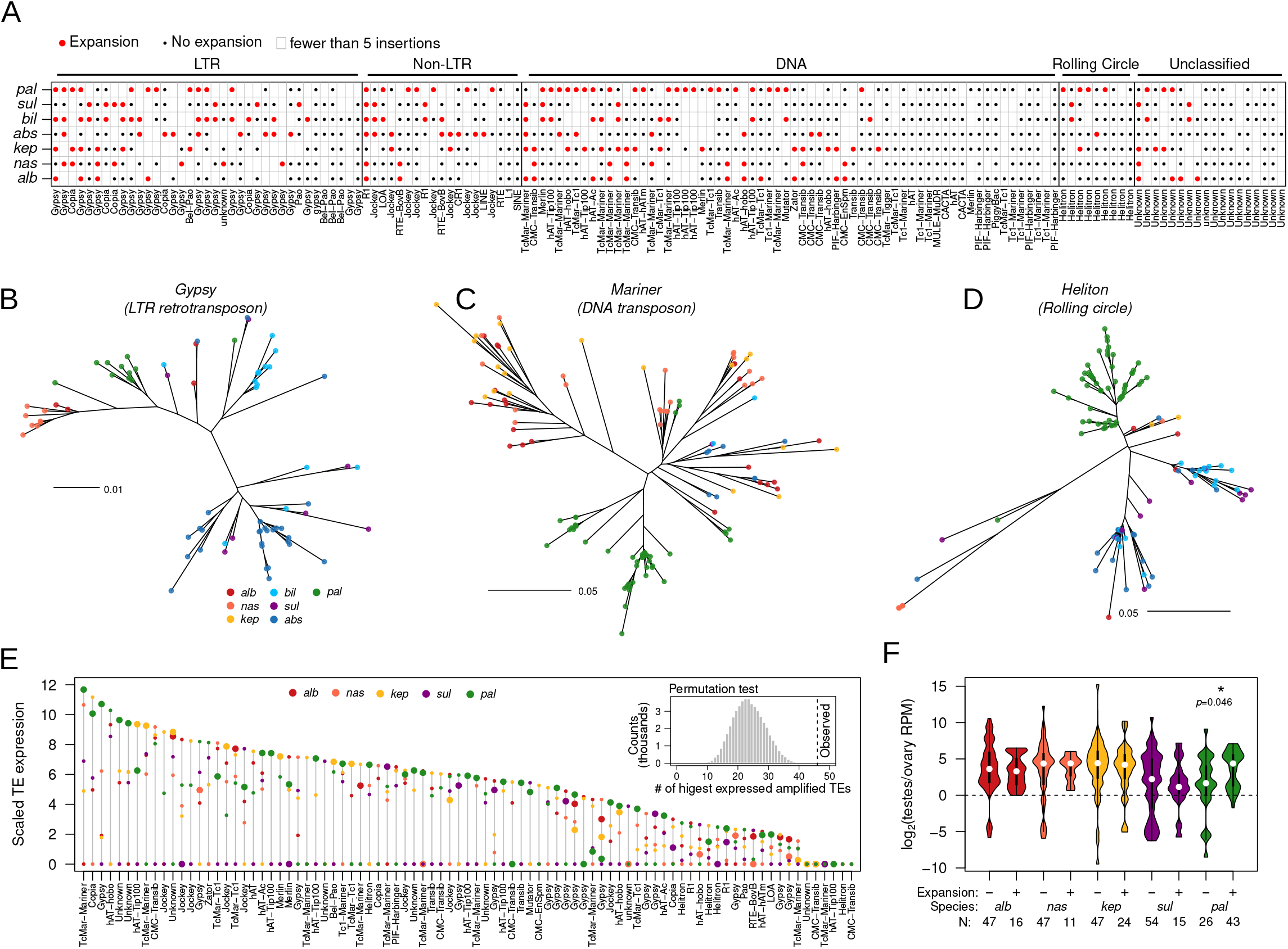
Frequent lineage specific amplifications and suppressions of TE families. A. Species-specific expansion status of different TE families and types based on phylogenies of insertions. Red dots indicate amplification in a *nasuta* species, black dots indicate no amplification, and empty boxes indicate fewer than 5 insertions. B-D. unrooted trees of TE insertions of different types of TEs. Their positions on the table in A are marked by arrowheads. E. Expression of expanded and unexpanded TE families in the testes of different species. For each TE family, the transcript abundance is scaled by the lowest expressed species, and the range of expression across the different species is plotted vertically as demarcated by the gray line. Along this line the expression in the different species are positioned by colored circles. Large circles denote species-specific expansion. The observed positions of the expanded TEs along the expression ranges are tested against the null expectation using randomized permutation testing (top right inset). The null distribution is presented and the observed count is marked by the vertical dotted line. F. Fold-difference in TE transcript abundance between testes and ovarian expression across species. TEs are subdivided into those that have species-specific expansions and those without.

To determine whether these expansions resulted from escape of transcriptional and post-transcriptional silencing, we examine TE expression from the testes and ovaries in five species. Cross-species comparisons revealed that TEs frequently show elevated expression accompanying their expansion (**Figure 6E**). Out of those that have expanded, 46 TE families (58.9%) show the highest expression in the species in which the expansion occurred, significantly higher than the random expectation of 24 (**Figure 6E**; p < 0.00002, permutation testing, see Materials and Methods). However, this is not always the case; for example, while DINE shows recurrent and recent expansions in *D. pallidifrons* (**Figure 5A**), it is expressed at intermediate levels in this species (**Figure S16**). Interestingly, we also find at least 15 instances where the TE family is the most lowly expressed in the species in which it expanded; we suspect these may reflect successful suppression mechanisms that evolved after expansion.

In Drosophila, the activity of TEs and their silencing systems can both differ between the sexes (Chen et al. 2021). Across all species, TE expression in testes is higher than in ovaries, suggesting weaker silencing in the testes. Curiously, expression of expanded TEs in *D. pallidifrons* are on average 20.70-fold higher in testes compared to ovaries. This is significantly higher than unexpanded TEs which are only 3.16-fold higher in the testes (p = 0.0464). This striking difference suggests that the numerous TEs that have expanded in *D. pallidifrons* may be exploiting the male germline for amplification which is consistent with our observation that insertions are found more frequently around genes with higher expression in testes compared to ovaries (**Figure 3C**).

### Epigenetic silencing of expanded TEs moderately reduces expression in neighboring genes

Even though expanded TEs are typically highly expressed when compared to other species, several expanded TEs show low to no expression. We hypothesized that the lowly expressed expanded TEs may have been historically active elements that are now silenced. To evaluate this possibility, we looked at expression of genes neighboring these expanded TEs, reasoning that silencing of TEs will likely lead to reduced expression of neighboring genes.

We identified genes with nearby TE insertions (internal or +/− 1kb up- and downstream), and subdivided them into those with insertions of highly vs. lowly expressed expanded TEs. We focused on *D. pallidifrons* as it has the highest number of expanded TEs, and identified 182 and 552 genes with expanded lowly expressed and expanded highly expressed nearby TEs, respectively. Interestingly, the expression of the former set (genes nearby highly expressed expanded TEs) are significantly higher than those of the latter (genes nearby lowly expressed expanded TEs; **Figure 7A**, p-value < 3.5392 × 10^−16^, Wilcoxon Rank Sum Test). This is consistent with the notion that silencing of expanded TEs is associated with lower expression of nearby genes.

**Figure 7.**
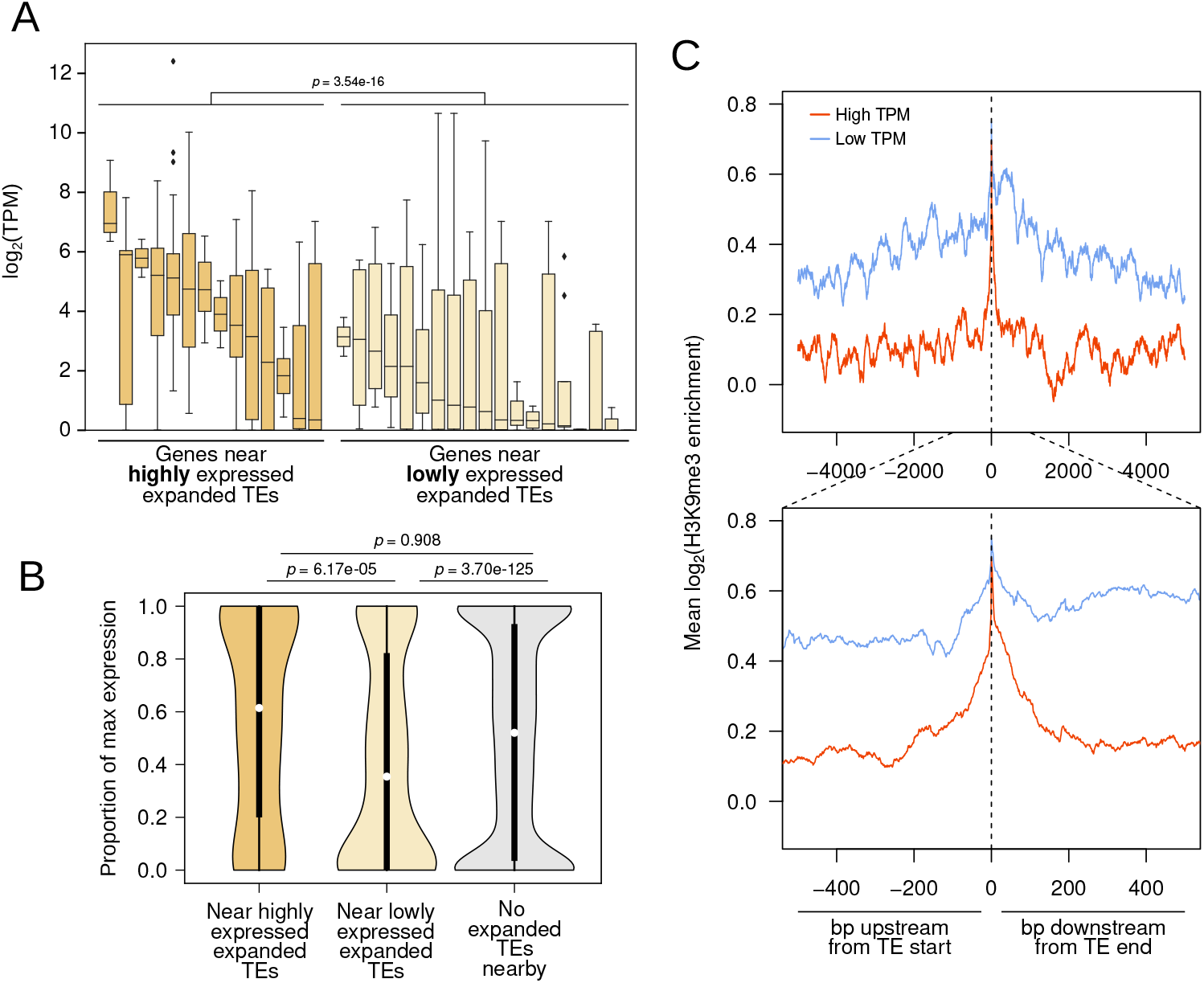
Epigenetic silencing of expanded TEs downregulates nearby genes. A. Expanded TEs are categorized as either highly or lowly expressed depending on expression difference between species. Dark yellow boxes represent genes nearby highly expressed expanded TEs, while light yellow boxes represent genes nearby lowly expressed expanded TEs. Each box represents the distribution of transcript abundances (TPM) of genes with nearby insertions of a given expanded TE family. Genes (n=785) near lowly expressed expanded TEs have significantly lower expression (Wilcoxon’s Rank Sum Test, p < 3.54e-16). B. Scaled expression of genes near highly (dark yellow, n=82) and lowly expressed expanded TEs (light yellow, n=552), as well as those with no expanded TEs nearby (gray, n=8705). Genic expression is scaled by the TPM of the highest expressed orthologs across all species. Significance of pairwise comparisons of the three sets are labeled above the figure. C. H3K9me3 enrichment around sull length TE insertions in D. albomicans depending on whether the TE is highly expressed as compared to other species (red) vs lowly expressed (blue). Insertions within the pericentric regions are removed.

To differentiate between insertions/survival bias near lowly expressed genes versus bona fide spreading of epigenetic silencing into neighboring genes, we again compared the expression of the orthologs of these genes between species. To sensitively detect potential down regulation, for each gene, we scaled the expression of the *D. pallidifrons* ortholog relative to the most highly expressed ortholog. For genes with no expanded TEs around them (**Figure 7B**, gray), the *D. pallidifrons* orthologs, expectedly, have a median relative expression of 0.50. Although not significantly different, genes with highly expressed expanded TEs nearby show a slightly higher median expression and are slightly skewed towards higher expression (**Figure 7B**, dark yellow). On the other hand, genes near lowly expressed expanded TEs (i.e. near those TEs that are putatively silenced) show a low relative expression of 0.37 (**Figure 7B**, light yellow). These genes show a clear skew towards low to no expression, and are significantly lower ranked than both the control set of genes (no expanded TEs nearby) and genes near highly expressed TEs (**Figure 7B**, light yellow; p= 3.70e-12 and 5.17e-05, Wilcoxon’s Rank Sum Test). These results reveal that insertions of recently expanded TEs can cause a subtle but significant decrease in gene expression if inserted nearby, but only if the TEs are targeted for (presumably epigenetic) silencing. However, if a recently expanded TE is not being targeted for silencing, it may potentially induce higher expression of neighboring genes.

We used our H3K9me3 ChIP data in *D. albomicans* to further evaluate whether this effect is due to epigenetic silencing. We plotted H3K9me3 enrichment around TEs with elevated expression and TEs with low expression in *D. albomicans*, removing insertions in the pericentric regions (**Figure 7C**). Consistent with epigenetic spreading at putatively silenced TEs, we find that TEs with low expression show substantially higher H3K9me3 enrichment in surrounding regions, with both elevated and wider spreading of heterochromatin. More highly expressed TEs, in contrast, show substantially less enrichment and spreading of H3K9me3. Therefore, lowly expressed TEs are likely under stronger epigenetic silencing which leads to broader spreading of H3K9me3.

## Discussion

Here, we generated repeat-rich genomes of seven closely related Drosophila species, taking advantage of long read sequencing technologies. Enabled by these high quality genome assemblies, we systematically characterized the landscape of TE insertions and evaluated how their activities and regulation influence genome evolution. Specifically, we focused on two questions: how often do TEs influence gene regulation and how common do TEs escape silencing and expand in copy number?

### The regulatory impact of TE insertions on gene expression

There are numerous examples of TE insertions affecting expression of neighboring genes, some of which even confer adaptive phenotypes (Merenciano et al. 2016; Casacuberta and González 2013; Villanueva-Cañas et al. 2019; Mateo et al. 2014). However, insertions around genes are primarily thought to be deleterious as they can induce epigenetic silencing of neighboring genes through heterochromatin spreading (Choi and Lee 2020). Here we comprehensively evaluate such an effect in a comparative genomics framework by combining high confidence TE insertion calls from de novo genome assemblies with gene expression data across a group of recently diverged species, the *nasuta* species group. While TE insertions are found more frequently near lowly expressed genes, TE-induced silencing does not appear to be a major cause of this negative association. The vast majority of genes with insertions around them do not show lower expression compared to other species. Therefore, instead of TEs causing nearby down-regulation, it appears that they tend to accumulate and repeatedly insert near historically lowly expressed genes. The fact that independent insertion patterns are positively correlated between species suggest that two types of non-mutually exclusive biases could be at play. TEs may preferentially insert into specific regions, chromatin environments, or gene features resulting in similar insertion patterns between species. This alone is unlikely to fully account for the negative association between gene expression and insertion counts. We, therefore, suspect that the observed insertion landscape also reflects a survivorship bias; insertions with high fitness costs are unlikely to reach high population frequency, therefore most of the observable insertions in the genome will be those with low fitness impacts. Unlike highly expressed genes, such as housekeeping genes that are under strong negative selection, lowly expressed genes may be more permissive to fluctuations in gene expression.

TEs are underrepresented near highly expressed genes, yet most TE insertions identified in our genomes do not appear to alter gene expression (**Figures 2F** and **3G**). If the observed TE insertions rarely influence gene expression, then how could they be more deleterious when inserted near highly expressed genes? One possible solution to this apparent paradox may be that the regulatory effects of TEs become more substantial upon environmental perturbations (Capy et al. 2000). In plants, multiple classes of retrotransposons are activated upon stresses (Wessler 1996; Grandbastien et al. 1997), and in flies and worms, TEs increase in activity during elevated temperatures (Garza et al. 1991; Ratner et al. 1992; Kidwell et al. 1977; Kurhanewicz et al. 2020). The lack of expression change in genes with TEs inserted nearby may therefore be the product of maintaining stocks in stable lab conditions. But upon environmental perturbation, these genes might begin to show more drastic regulatory changes as TEs become active. In changing environmental conditions, insertions around highly expressed and functionally important genes may therefore be under strong negative selection, accounting for the negative association between gene expression and insertions.

### Context dependent heterochromatin spreading and epigenetic silencing

Despite no systematic support for widespread downregulation of genes with TEs inserted nearby, we were able to find evidence of epigenetic silencing of genes due to insertions in some cases. In *D. pallidifrons*, insertions of recently expanded TEs can cause moderate down regulation of gene expression, but only if the TEs have low expression - presumably due to epigenetic silencing (**Figure 7A** and **B**). Moreover, in every species a few dozens of species-specific TE insertions appear to be associated with down-regulation of nearby genes (**Figure 3G** and **H**, **Table S4**). Notably, we also find cases where insertions are associated with large up-regulation in gene expression (**Figure 3G** and **H**, **Table S4**), but these cases are much rarer than those associated with down-regulation of nearby genes.

While we do not have direct evidence of transcriptional or post-transcriptional silencing in most species, clear spreading of heterochromatin from TE insertions is observed in *D. albomicans* (**Figure 4**). TEs far from genes show the highest and broadest H3K9me3 enrichment, and TE insertions near lowly expressed genes also show more heterochromatin spreading. While consistent with epigenetic silencing of neighboring genes, these results are also consistent with the notion that active transcription antagonizes heterochromatin formation, and vice versa. Lower expression of genes near TEs that show higher levels of heterochromatin spreading could indicate that H3K9me3-inducing TEs are more tolerated near lowly expressed genes. Further, we find that insertions of TE families with low expression are associated with broader and stronger heterochromatin spreading to their surroundings. Indeed, lowly expressed and high copy number TEs are typically recently active and have robust small RNA targeting for post-transcriptional degradation and transcriptional silencing (Wei et al. 2021). Altogether, these results suggest that TE insertions can have multiple effects on gene expression and calls into question how pervasive TE-induced epigenetic silencing of neighboring genes is. The epigenetic effects TEs have on neighboring genes, if any, is likely dependent on multiple factors, such as the transcription rate of the gene, the local repeat density and the 3D architecture of the genome.

### Frequent and recurrent TE expansions and silencing

Using a phylogenetic approach to understand the relationship of TE insertions, we revealed that >50% of the TE families show lineage and species-specific amplification. The most striking expansion is the DINE, which has exploded to thousands of copies across the *nasuta* species group. This expansion occurred once prior to the species radiation, and at least twice since, one in *D. pallidifrons*, and one in the related outgroup species *D. immigrans* (~20 million years diverged; Izumitani et al. 2016; O’Grady and DeSalle 2018) (**Figure 5A**). The repeated expansions suggest multiple bouts of suppression escape. Interestingly, we were unable to find unique mutations private to the *D. pallidifrons* expansion clade, which may be causal mutations allowing to avoid suppression. One possible explanation for the absence of such mutations is that gene conversion events have converted some of such nucleotides in insertions within the clade to nucleotides from other variants and vice versa, causing more polymorphic distribution of the nucleotides (Fawcett and Innan 2019). Such events have previously been shown to allow rapid adaptive changes at TE sequences co-opted for X-chromosome dosage compensation (Ellison and Bachtrog 2015). Consistent with gene conversion, there are multiple smaller scale clusters of *D. pallidifrons* DINEs all across the tree which may represent elements that acquired the causal mutations allowing for their own, albeit limited, suppression escapes. Previous analyses of DINEs across *Drosophila* have found their sequences to be species-specific (Yang and Barbash 2008), even for recently diverged species. This may be due to rapid homogenization of copies due to gene conversion events similar to what we are observing in *D. pallidifrons*.

Beyond DINEs, large fractions of TEs also show lineage specific expansions, though at much more limited scales. Most of these expanding TE families show elevated expression only in the species with the expansion, consistent with species-specific suppression escape and derepression allowing for expansion. Most strikingly, 32 families are or have been recently expanding in *D. pallidifrons*. This may in part reflect the fact that it is the least derived of our species and therefore has the longest terminal branches. However, we still find high expression for many of these expanding TEs indicating recent, and perhaps, on-going mobilizations. Why are so many TEs concurrently expanding in *D. pallidifrons*? P-element dysgenesis is caused by the absence of maternally deposited piRNAs against the P-elements, yet derepression and mobilization of TEs is not limited to P-elements (Khurana et al. 2011). Therefore, the large numbers of expanding and highly expressed TEs may be reflecting an on-going sweep of a novel TE in the species. Interestingly, we also find that a fraction of these recently expanded TEs, paradoxically, have low expression, and genes around them show reduced expression. We suspect that these are recently active TEs that are now epigenetically silenced.

The importance of horizontal transfer to the long-term survival and expansion of TEs has been pointed out multiple times in the literature (Kidwell 1992; Silva et al. 2004; Loreto et al. 2008; Schaack et al. 2010; Zhang et al. 2020). Horizontal transfer can allow TEs to cross species-boundaries and invade a naive genome that lacks suppressive mechanisms against this TE, where it can proliferate (Le Rouzic and Capy 2005). Once silencing mechanisms against a TE are in place, for example targeting by small RNAs, mobilization of that TE is prevented (Khurana et al. 2011). Inactive TEs will accumulate mutations, and eventually all functional copies may die, and horizontal transfer to a new lineage would allow that TE to escape extinction. Our finding of species-specific escape from TE repression for a large fraction of TE families suggests a very dynamic evolution of host genomes and their TEs. Active TEs are temporarily silenced within a lineage, but over evolutionary timescales, some copies will escape silencing in different lineages, leading to species-specific bursts in TE activity. Thus, in addition to horizontal transfer, our data suggest that escape from host suppression seems to be an important strategy allowing for the long-term survival of TEs.

### Long-read genome assemblies open new doors for studying TEs

In our study, high quality genomes assembled via long reads have circumvented many of the previous challenges associated with studying TEs and repeats (Khost et al. 2017), and enabled high confidence annotation of TE insertions. Further, our approach of integrating phylogenetics, functional genomics, and comparative genomics have revealed a comprehensive picture of the dynamics of TE suppression escape and subsequent re-established silencing and their effects on the rest of the genome. These high quality genome assemblies will further facilitate the molecular dissection of the nucleotide changes in TEs causing suppression escape in future studies. With the rapidly decreasing cost and input material in generating these assemblies (Adams et al. 2020), it will become easier and cheaper to identify de novo insertions. But even with the rapid adoption of these technologies, TEs and repeats remain under-studied and often avoided. Instead, here we show that assembling repeats is among one of the greatest advantages to long read sequencing.

## Material and Methods

### Fly strains and nanopore sequencing

We extracted high molecular weight DNA from approximately 50 females from *D. nasuta* 15112-1781.00, *D. kepulauana* 15112-1761.03*, D. s. albostrigata* 15112-1771.04*, D. s. bilimbata* 15112-1821.10*, D. s. sulfurigaster* 15112-1831.01, and *D. pallidifrons* PN175_E-19901 using the QIAGEN Gentra Puregene Tissue Kit. The *D. kepulauana* high molecular weight DNA was sequenced on PacBio RS II platform at UC Berkley QB3 genome sequencing center.The high molecular weight DNA of other species were sequenced on Nanopore MinIOn.

### Genome assemblies

#### Drosophila albomicans

The *D. albomicans* genome assembly has been previously published, having been generated with DNA sequenced on the PacBio RSII platform resulting in an N50 of 33.4 Mb and BUSCO score of 98%, indicating high contiguity and completeness. Here, Nanopore reads from strain 15112-1751.03 were error corrected with canu and an initial assembly was generated using wtdbg2. The assembly was then polished 3 times using 35.7x coverage Illumina paired end reads from *Mai et al.* 2019 with minimap2 and Racon followed by 1 round of Pilon. Afterwards, we BLAST the assembly against the NCBI BLAST database for potential contamination. We remove 65 contigs making up 1.4 Mb—mostly comprising *Acetobacter*. This filtered genome is then organized using HiC data and the Juicer and 3d-dna pipeline. We stitch adjacent contigs within a scaffold with a string of 50 N’s. This stitched genome assembly has an N50 of 33,438,794 bp and a BUSCO score of 91.3%, notably lower than the previously published assembly. To improve upon this assembly, we use quickmerge twice with these two assemblies and twice more with the results, taking complementary information between them to improve contiguity and completeness. Due to the reference dependent asymmetry of quickmerge results, we ran the program using both the old and newly stitched genome as the reference and repeated this with the resulting genomes; we took the one with the highest contiguity and BUSCO score. We generated an even more complete assembly (BUSCO score of 99.6%), an increase in assembly size (167,541,436 bp) and improved contiguity (N50 of 35,291,776 bp).

#### Drosophila nasuta

The *D. nasuta* genome assembly was generated using Nanopore long read data from strain 15112-1781.00, which were error corrected with canu. The initial genome assembly was generated using wtdbg2 and polished—using 38.3x coverage Illumina paired end reads from *Mai et al.* 2019 3 times with Racon and minimap2 followed by 1 round of Pilon. We BLAST the assembly against the NCBI database for contamination and find 140 contigs making up approximately 5.57 Mb, mostly from *Acetobacter*. This filtered genome is organized with HiC data using the Juicer and 3d-dna pipeline, where we stitched adjacent contigs with a string of 50 N’s. The final resulting assembly has a BUSCO score of 99.2%, assembly size of 171,781,232 bp, and an N50 of 33,885,645 bp.

#### Drosophila kepulauana

The *D. kepulauana* genome assembly was generated with DNA sequenced on the PacBio RSII platform data from strain 15112-1761.03—the reads were error corrected with canu. Similar to the *D.* albomicans assembly, we generated two genomes and used quickmerge to generate the final assembly. The first assembly was initially generated using wtdbg2 and polished with 38x coverage Illumina paired end reads from *Mai et al.* 2019 thrice with Racon and minimap2 followed by 1 round of Pilon. We BLAST the assembly against the NCBI database for contamination and find 87 contigs making up approximately 14.03 Mb, mostly from *Acetobacter*. This filtered genome is organized with HiC data using the Juicer and 3d-dna pipeline, where we stitched adjacent contigs with a string of 50 N’s. The second assembly was initially generated with Flye. This assembly was polished and filtered for contamination (58 contigs totaling 13.29 Mb) in the same way as the first assembly. We ran quickmerge twice on the two assemblies, using each one as the reference, and twice more on the resulting assemblies. The assembly deemed as the final assembly is the most contiguous and complete one and has a BUSCO score of 99.7%, assembly size of 163,769,021 bp, and an N50 of 34,564,094 bp.

#### Drosophila sulfurigaster albostrigata

The *D. s. albostrigata* genome assembly was generated using nanopore sequencing data from strain 15112-1771.04, which were error corrected with canu. Just like the treatment of the *D. kepulauana* assembly, we generated two genomes and used quickmerge to generate the final assembly. The first assembly was initially generated using wtdbg2 and polished with 39.4x coverage Illumina paired end reads from *Mai et al.* 2019 Illumina paired end short reads three times with Racon and minimap2 followed by 1 round of Pilon. We BLAST the assembly against the NCBI database for contamination and find 8 contigs making up approximately 7.1 Mb, mainly from *Acetobacter*. This filtered genome is ordered with HiC data using the Juicer and 3d-dna pipeline and adjacent contigs were stitched with a string of 50 N’s. The second assembly was initially generated with Flye. This assembly was polished and filtered for contamination (25 contigs totaling 7.71 Mb) in the same way as the first assembly. We ran quickmerge twice on the two assemblies, using each one as the reference, and twice more on the resulting assemblies. The assembly deemed as the final assembly is the most contiguous and complete one and has a BUSCO score of 98.5%, assembly size of 168,284,230 bp, and an N50 of 37,627,869 bp.

#### Drosophila sulfurigaster bilimbata

The *D. s. bilimbata* genome assembly was generated using Nanopore long read data from strain 15112-1821.10, which were error corrected with canu. The initial genome assembly was generated using wtdbg2 and polished using 17.1x coverage Illumina single end reads from *Mai et al.* 2019 Illumina single end short reads 3 times with Racon and minimap2 followed by 1 round of Pilon. We BLAST the assembly against the NCBI database for contamination and find 186 contigs totaling around 7.41 Mb, mostly from *Acetobacter*. This filtered genome is organized with HiC data using the Juicer and 3d-dna pipeline, where we stitched adjacent contigs with a string of 50 N’s. The final resulting assembly has a BUSCO score of 99.7%, assembly size of 164,595,183 bp, and an N50 of 36,279,119 bp.

#### Drosophila sulfurigaster sulfurigaster

The *D. s. sulfurigaster* genome assembly was generated using Nanopore long read data from strain 15112-1831.01, which were error corrected with canu. An initial genome assembly was generated using Flye and polished—using 26.7x coverage Illumina paired end reads from *Mai et al.* 2019--3 times with Racon and minimap2 followed by 1 round of Pilon. We BLAST the assembly against the NCBI database for contamination and find 11 contigs totaling around 3.3 Mb, most of which were from *Acetobacter*. This filtered genome is ordered with HiC data using the Juicer and 3d-dna pipeline, where we stitched adjacent contigs with a string of 50 N’s. The final resulting assembly has a BUSCO score of 99.9%, assembly size of 165,884,258 bp, and an N50 of 35,818,991 bp.

#### Drosophila pallidifrons

The *D. pallidifrons* genome assembly was generated using Nanopore long read data from strain PN175_E-19901, which were error corrected with canu. The initial genome assembly was generated using wtdbg2 and polished—using 17x coverage Illumina paired end reads from *Mai et al.* 2019 3 times with Racon and minimap2 followed by 1 round of Pilon. We BLAST the assembly against the NCBI database for contamination and find 59 contigs making up approximately 4.9 Mb, mostly from *Acetobacter*. This filtered genome is organized with HiC data using the Juicer and 3d-dna pipeline, where we stitched adjacent contigs with a string of 50 N’s. The final resulting assembly has a BUSCO score of 99.3%, assembly size of 164,659,715 bp, and an N50 of 37,973,042 bp.

### Gene annotation and clustering

We used MAKER to annotate genes in each species’ genome assembly (Campbell et al. 2014). To train MAKER’s gene inference model, we generated a transcriptome from *D. albomicans* and *D. nasuta* RNA seq data from Zhou and Bachtrog 2012 (Zhou and Bachtrog 2012). RNA seq data from *D. albomicans* and *D. nasuta* were aligned to the corresponding genome assemblies with HISAT2 under default settings (Kim et al. 2019). The alignments were then used to create transcriptomes using StringTie (Pertea et al. 2015). Additionally, satellite repeats in the genome assemblies for each species were masked using RepeatMasker in preparation for gene annotations([CSL STYLE ERROR: reference with no printed form.]). Then, using both the *D. albomicans* and *D. nasuta* transcriptomes, we ran MAKER with default settings. We then took the annotations and determined gene homology between species with OrthoDB (Kriventseva et al. 2019).

### TE library generation, annotation and analyses

In order to lower the occurrence of nested TE structures, pericentromeric regions were removed from each and the resulting sequences were separately used as input for RepeatModeler2 and the accompanying LTRharvest software with default options (Flynn et al. 2020; Ellinghaus et al. 2008). The resulting species-specific TE libraries were merged together. To remove redundancy from the merged library, we used CD-hit2 to cluster TE entries with each. However, instead of allowing CD-hit2 to select the representative sequence of the cluster (which is usually the longest sequence), we evaluated the TEs within clusters based on three criteria: entry sequence length, self-identity, and probability of full length insertions. For self-identity, we blasted each TE entry to itself and calculated the self-blast score as the proportion of the sequence showing alignment to another region of itself. For probability of full length insertions, we blasted each entry to the genome and proportion of near full length blast hits. We then weighed the three criteria to maximize length and probability of full length insertion, while minimizing self-identity, in order to select the representative sequence per CD-hit2 cluster. This procedure is done twice. We used both the Repeatmodeler2 TEs categorization as well as the program ClassifyTE (Panta et al. 2021). When the two disagreed with the TE classification, we used the assignment from RepeatModeler2. Note, even after two round of CD-hit2 we found 10 redundant entries corresponding to variants of the DINE in the genome through manual NCBI BLASTn (Altschul et al. 1990). We removed entries with unique sequences flanking the DINEs and kept the longest entry.

TE insertions in genomes are annotated by RepeatMasking the final nasuta group-specific TE index to the respective species genomes. Because RepeatMasker can provide overlapping annotations, we used bedtools merge to merge overlapping annotations first, generating chimerics. We then blasted all the chimeric annotations to the repeat library and recategorized those where 90% of the sequence blasts to a specific TE. Full length and truncated elements are defined as annotations that are >80% length of the TE entries, or <80% length but >200 bp, respectively. Distances between full length TE and the closest gene in each species were calculated using bedtools closest, (Quinlan and Hall 2010) with species-specific TE and gene annotations as inputs.

### Phylogenetic analyses of TEs

We ran BLAST using the TE libraries as the query and the genome assemblies of each species as the database (Altschul et al. 1990). TE sequences from full length BLAST alignments--defined as those in which the alignment length is at least 80% of the TE length--are extracted. We used Clustal Omega under default settings to perform a multiple sequence alignment for all sequences for each TE (Sievers et al. 2011); those with over 200 full length copies across all species were subsampled down to 200 sequences. In order to maintain the different copy number in the different species, the subsampling procedure maintained the proportional difference of insertion counts across the species.

Phylogenies for TEs were then generated with RAxML using the command: raxmlHPC-PTHREADS-AVX -T 24 -f a -x 1255 -p 555 -# 100 -m GTRGAMMA -s input.MSA.fa -n input.MSA.tree > input.MSA.tree.stderr (Stamatakis 2014).

We tested for the presence of species specific expansion of each TE by measuring the extent of clustering using the RRphylo R package (Serio et al. 2019). Tests were carried out for species where there were at least 5 sequences or 5% of the total sequences in the phylogeny. The resulting p-values from the analyses were adjusted for multiple testing using the Benjamini-Hochberg procedure. TEs from a particular species with p-values < 0.05 are considered to be expanded. We note that this program does not take into account of the species relationships and therefore cannot capture lineage-specific expansion. Thus this approach under-estimates the number of TEs that have recently expanded.

### RNA sample collection and sequencing

Two replicates of RNA sequencing libraries created from males and females of each species were generated and sequenced. Testes from five to eight males from live *D. albomicans, D. nasuta, D. kepulauana,* and *D. s. sulfurigaster* as well as frozen *D. pallidifrons* were dissected for each RNA sequencing library. Ovaries from three to five females from live *D. albomicans, D. nasuta, D. kepulauana,* and *D. s. sulfurigaster* as well as frozen *D. pallidifrons* were dissected for each RNA sequencing library. For each species, tissue samples were placed in Trizol for RNA extraction. RNA was extracted using the Trizol extraction method and enriched for ployA RNA using NEBNext Poly(A) mRNA Magnetic Isolation Module (E7490) as per manufacturer protocol. The RNA libraries prepared as per NEBNext Ultra II Directional RNA kit (E7760S) and sequenced on illumina NovSeq 6000 on SP flow cell for 150 PE reads.

### RNA transcript abundance

#### Genes

Generated RNA sequencing data for *D. albomicans, D. nasuta, D. kepulauana, D. s. sulfurigaster,* and *D. pallidifrons* were aligned to their corresponding genome assembly. Using the alignment data and gene annotations, we used the featureCounts program from the Subread package to calculate the number of reads mapping to each gene. We then calculated gene transcripts per million with the following formula:

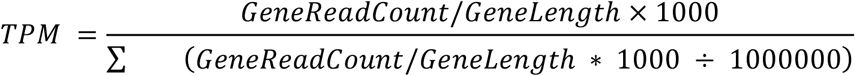

#### Transposable Elements

Generated RNA sequencing data for *D. albomicans, D. nasuta, D. kepulauana, D. s. sulfurigaster,* and *D. pallidifrons* were aligned to the TE library. A custom script was used to count the number of reads mapping to each transposable element. The number of reads was then normalized by the TE length and then divided by the median of gene read counts that are normalized in the same way from the corresponding species.

### Permutation testing of TE expression

The test statistic used for the permutation test is the number of times the highest expression for a particular TE comes from a species where that TE has expanded. We first calculate this test statistic from our data. We then randomly shuffle the species associated with each expansion event and calculate the test statistic 50,000 times. The p-value obtained is the proportion of tests with test statistics less than or equal to our original test statistic.

### TE expression comparisons

We categorize whether TEs are highly expressed or lowly expressed upon obtaining normalized TE expression level for testes and ovaries across species. A TE is considered to be highly expressed in a species for a particular tissue if the expression level of the TE is in the top two most highly expressed TE across species in that tissue. A TE is considered to be lowly expressed if it, instead, is in the bottom two most lowly expressed TE across species.

To compare expression between highly expressed TEs and lowly expressed TEs within a species, we first scale a TE’s log_2_ expression to its maximum log_2_ expression across species:

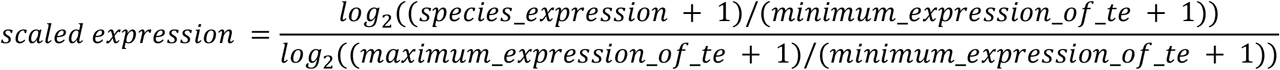

The addition of 1 to the values are done to prevent potential division by zero. We then perform the Wilcoxon rank sum test between scaled expression of highly expressed TEs and scaled expression of lowly expressed TE to determine if scaled expression between TEs from different categories are statistically significant.

### ChIP-seq analyses

ChIP-seq analyses were slightly modified from methods in (Wei et al. 2021). Briefly, larval H3K9me3 ChIP and input data (Wei and Bachtrog 2019) were aligned to the genome using bwa mem. The per base pair coverage was determined using bedtools coverageBed -d -ibam. Median autosomal coverage was estimated in from 50kb non-overlapping sliding windows. We then inferred enrichment at every position as:

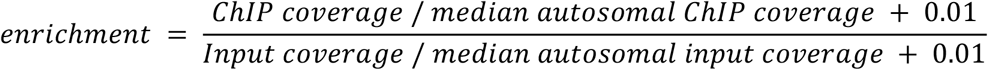

We averaged the enrichment across the three replicates. For H3K9me3 spreading around TE insertions, we lined up annotated TE insertions at either the 5’ or 3’, and averaged enrichment 5kb upstream and downstream of the insertions, respectively.

